# Accurate measurement of the effects of all amino-acid mutations to influenza hemagglutinin

**DOI:** 10.1101/047571

**Authors:** Michael B. Doud, Jesse D. Bloom

## Abstract

Influenza genes evolve mostly via point mutations, and so knowing the effect of every amino-acid mutation provides information about evolutionary paths available to the virus. We previously used high-throughput mutagenesis and deep sequencing to estimate the effects of all mutations to an H1 influenza hemagglutinin on viral replication in cell culture (Thyagarajan and Bloom, 2014); however, these measurements suffered from substantial noise. Here we describe advances that greatly improve the accuracy and reproducibility of our measurements. The largest improvements come from using a helper virus to reduce bottlenecks when generating viruses from plasmids. Our measurements confirm that antigenic sites on the globular head of hemagglutinin are highly tolerant of mutations. However, other regions – including stalk epitopes targeted by broadly neutralizing antibodies – have a limited capacity to evolve. The ability to accurately measure the effects of all influenza mutations should enhance efforts to understand and predict viral evolution.

## Introduction

Seasonal influenza is a recurrent threat to human health largely because it rapidly accumulates amino-acid mutations in proteins targeted by the immune system (Smith et al., 2004). Measuring the functional impact of every possible amino-acid mutation to influenza can therefore provide useful information about which evolutionary paths are accessible to the virus. Such measurements are now possible using deep mutational scanning (Fowler and Fields, 2014; Boucher et al., 2014). When applied to influenza, this technique involves creating all codon mutants of a viral gene, incorporating these mutant genes into viruses that are subjected to a functional selection, and estimating the functional impact of each mutation by using deep sequencing to quantify its frequency pre- and post-selection. We and others have used deep mutational scanning to estimate the effects of all amino-acid or nucleotide mutations to several influenza genes (Thyagarajan and Bloom, 2014; Bloom, 2014a; Wu et al., 2014; Doud et al., 2015; Wu et al., 2016). However, these studies suffered from substantial noise that degrades the utility of their results. For instance, in every study that reported the results for independent experimental replicates, the replicate-to-replicate correlation was mediocre.

This experimental noise arises primarily from bottlenecking of mutant diversity during the generation of viruses from plasmids. The influenza genome consists of eight negative-sense RNA segments. During viral infection, gene expression from these segments is a highly regulated process (Chua et al., 2013; Robb et al., 2009; Shapiro et al., 1987). Generating influenza from plasmids involves cotransfecting mammalian cells with multiple plasmids that must yield all eight viral gene segments and at least four viral proteins at a stoichiometry that leads to assembly of infectious virions (Hoffmann et al., 2000; Neumann et al., 1999; Fodor et al., 1999). This plasmid-driven process is understandably less efficient than viral infection. A small fraction of transfected cells probably yield most initial viruses, which are then amplified by secondary infection. This bottlenecking severely hampers experiments that require creating a diverse library of viruses from an initial library of plasmids.

Several strategies have been used to overcome problems associated with bottlenecks during the generation of influenza from plasmids. One strategy is to generate and titer each viral variant individually, and then mix them (Varble et al., 2014; Benitez et al., 2015). A second strategy is to reduce the impact of bottlenecks by shrinking the complexity of the libraries, such as by only mutating a small portion of a viral gene (Wu et al., 2015; Jiang et al., 2015). Neither of these strategies scale effectively to the deep mutational scanning of full-length proteins, since there are ~ 10^4^ unique amino-acid mutants of a 500-residue protein.

To overcome these limitations, we have developed a novel approach that uses a “helper virus” to generate virus libraries without strong bottlenecking. We have combined this approach with other technical improvements to repeat our deep mutational scanning of all amino-acid mutations to an H1 hemagglutinin (HA) with much higher accuracy and reproducibility. We use phylogenetic analyses to show that our new measurements better reflect constraints on HA evolution in nature. We confirm that antigenic sites in the globular head of HA are highly tolerant of mutations, and identify other regions of the protein that are more constrained. These advances improve our understanding of HA’s inherent evolutionary capacity, and can help inform evolutionary modeling and guide the development of vaccines targeting sites with a limited capacity for mutational escape.

## Results

### A helper-virus enables efficient production of mutant virus libraries from plasmids

We reasoned that the process of generating viral libraries carrying HA mutants would be more efficient if transfected cells only needed to produce HA from plasmid, and the other gene segments and proteins were delivered by viral infection (Figure 1A). Marsh et al. (2007) described seven-segmented HA-deficient virus that could be propagated in cells that constitutively express HA protein. We created HA-expressing cells and validated that we could propagate an HA-deficient A/WSN/1933 (H1N1) virus (Figure 1 - figure supplement 1).

**Figure 1.**
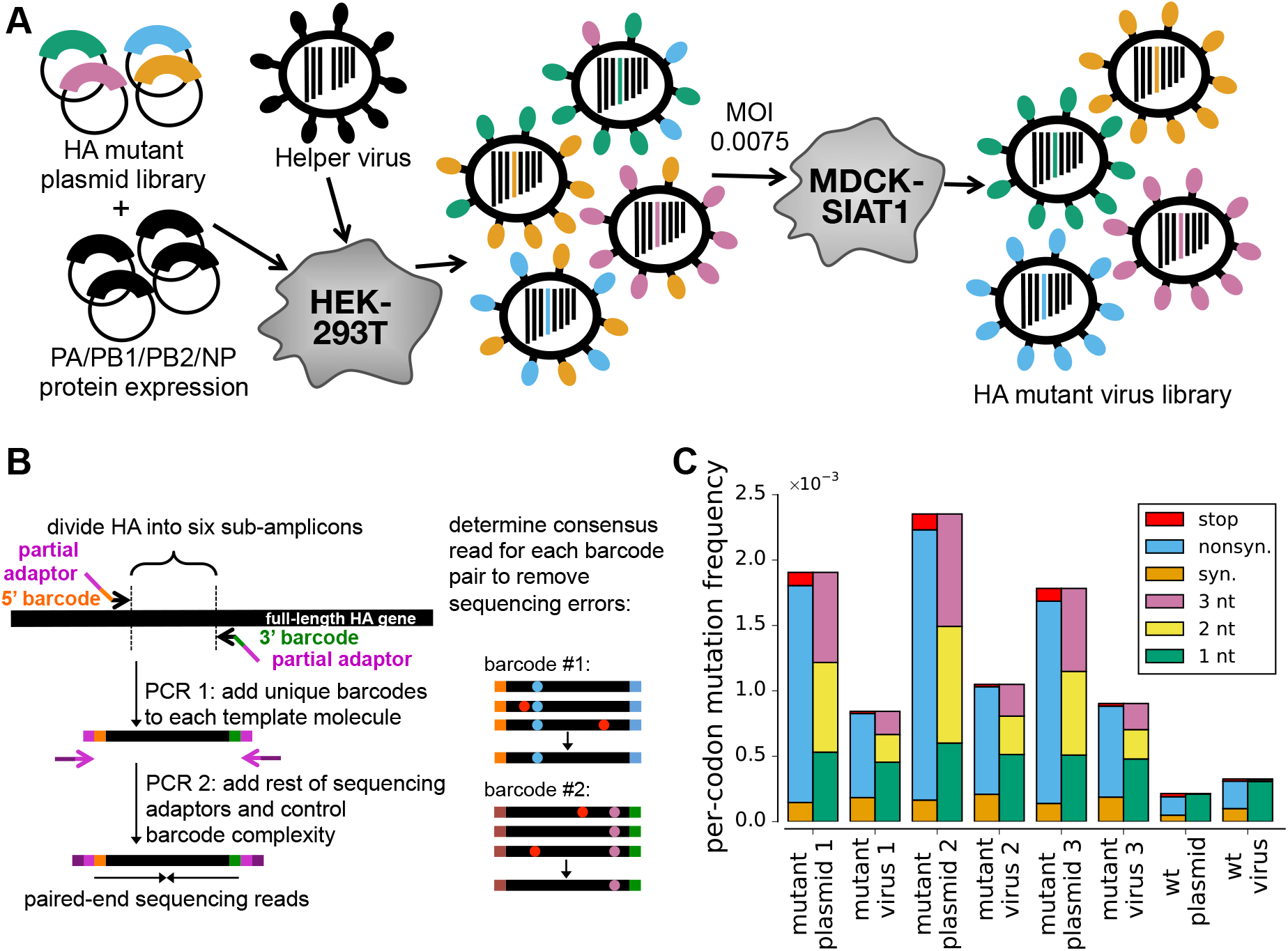
Deep mutational scanning of HA. **(A)** Cells transfected with a plasmid mutant library of HA are infected with an HA-deficient helper virus to yield a library of mutant viruses. This virus library is passaged at low MOI to select for functional variants and enforce genotype-phenotype linkage. The helper viruses themselves are propagated in cells constitutively expressing HA (Figure 1 - figure supplement 1). The variants in the plasmid mutant library contain an average of one codon mutation, with the number of mutations per clone following a roughly Poisson distribution (Figure 1 - figure supplement 2). The helper-virus works best when HA is provided on a plasmid that directs the synthesis of only viral RNA (Figure 1 - figure supplement 3). **(B)** Accurate Illumina sequencing using barcoded subamplicons. HA is divided into six sub-amplicons, and a first round of PCR appends random barcodes and part of the Illumina adaptor to each subamplicon. The complexity of these barcoded subamplicons is controlled to be less than the sequencing depth, and a second round of PCR adds the remaining adaptor. Sequencing reads are grouped by barcode, distinguishing sequencing errors that occur in only one read (red döts) from true mutations that occur in all reads (blue and purple döts). **(C)** The overall mutation frequencies reveal selection against stop codons and many nonsynonymous mutations in the mutant viruses relative to the plasmids from which they were generated (see also Figure 1 - figure supplement 4). Sequencing of unmutated plasmid and virus generated from this plasmid indicates tolerably low råtes of sequencing, reverse-transcription, and viral replication errors.

We cloned triplicate plasmid libraries of random codon mutants of the A/WSN/1933 HA gene. These libraries contain multi-nucleotide (e.g., `GGC→CAT`) as well as single-nucleotide (e.g., `GGC→GAC`) codon mutations. There are 63 × 565 ≈ 3.5 × 10^4^ different codon mutations that can be made to the 565-codon HA gene, corresponding to 19 × 565 ≈ 10^4^ amino-acid mutations. The deep sequencing described below found at least three occurrences of over 97% of these amino-acid mutations in each of the three replicate plasmid mutant libraries. These libraries have a somewhat lower mutation rate than those in Thyagarajan and Bloom (2014), with the number of mutations per clone following a roughly Poisson distribution with a mean of about one (Figure 1 - figure supplement 2). We cloned these HA libraries into both uni-directional and bidirectional reverse-genetics plasmids (Neumann et al., 1999; Hoffmann et al., 2000).

We then transfected cells with one of the HA plasmid mutant libraries along with plasmids expressing the four viral polymerase-related proteins (PB2, PB1, PA, and NP) with the goal of generating preformed viral ribonucleoprotein complexes carrying the HA segment. These transfected cells were then infected with HA-deficient helper virus, and 24 hours later we determined the titer of fully competent virus in the supernatant. The highest titers (∼ 10^3^ TCID_50_ per *µ*l) were obtained using the uni-directional reverse-genetics plasmid (Figure 1 – figure supplement 3). Overall, this result demonstrates the feasibility of the helper-virus strategy in Figure 1A.

We next used this helper-virus strategy to independently generate three mutant virus libraries, one from each of our triplicate plasmid mutant libraries. Each mutant virus library should sample most of the codon mutations to the A/WSN/1933 HA. We also generated a control virus library from a plasmid encoding the unmutated wild-type HA gene.

### Low MOI passage combined with barcoded-subamplicon sequencing reveals strong selection against non-functional HA variants

To select for viruses carrying functional HA variants, we passaged the mutant virus libraries at a low multiplicity of infection (MOI) of 0.0075 TCID_50_ per cell as outlined in Figure 1A. This MOI is substantially lower than that used in Thyagarajan and Bloom (2014) (0.1), and was chosen with the goal of more effectively purging non-functional HA variants.

To quantify selection on HA, we used deep sequencing to determine the frequency of each mutation pre- and post-selection. Standard Illumina sequencing has an error rate that is too high. In Thyagarajan and Bloom (2014), we reduced this error rate by using overlapping paired-end reads. Here we used an alternative error-correction strategy that involves attaching random barcodes to PCR subamplicons and then clustering reads with the same barcode (Figure 1B). To our knowledge, this basic strategy was first described by Hiatt et al. (2010) and first applied to influenza by Wu et al. (2014). Sequencing of the unmutated plasmid allows us to estimate that the error rate is ∼ 2 × 10^-4^ per codon, corresponding < 10^-4^ per nucleotide (Figure 1C). This error rate is substantially lower than we obtained previously using overlapping paired-end reads, consistent with the results of the sequencing-strategy comparison by Zhang et al. (2016). Sequencing of virus generated from the unmutated plasmid shows that the error rates associated with reverse-transcription and viral replication are also tolerably low (Figure 1C).

Figure 1C reveals strong selection against non-functional HA variants. The plasmid mutant libraries contain a mix of synonymous, nonsynonymous, and stop-codon mutations. However, stop-codon mutations are almost completely purged from the passaged mutant virus libraries, as are many nonsynonymous mutations. The selection against the stop codons is stronger than in Thyagarajan and Bloom (2014) (Figure 1 – figure supplement 4). Overall, these results indicate strong selection on HA that can be quantified by accurate deep sequencing.

### The mutant virus libraries have reduced bottlenecking and yield reproducible measurements of mutational effects

To evaluate whether the virus libraries were bottlenecked, we examined the distribution of synonymous mutation frequencies in each library. If bottlenecking causes a few mutants to stochastically dominate, we expect that in each library a few sites will have relatively high synonymous mutation frequencies and that these sites will differ among replicates. Figure 2A shows normalized synonymous mutation frequencies across HA for each of the three replicate mutant virus libraries from both Thyagarajan and Bloom (2014) and the current study. In the older study, each replicate has a different handful of sites with greatly elevated synonymous frequencies (green arrows), indicative of stochastic bottlenecking. In contrast, in our new virus libraries, the distribution of synonymous mutation frequencies is much more uniform across the HA gene.

**Figure 2.**
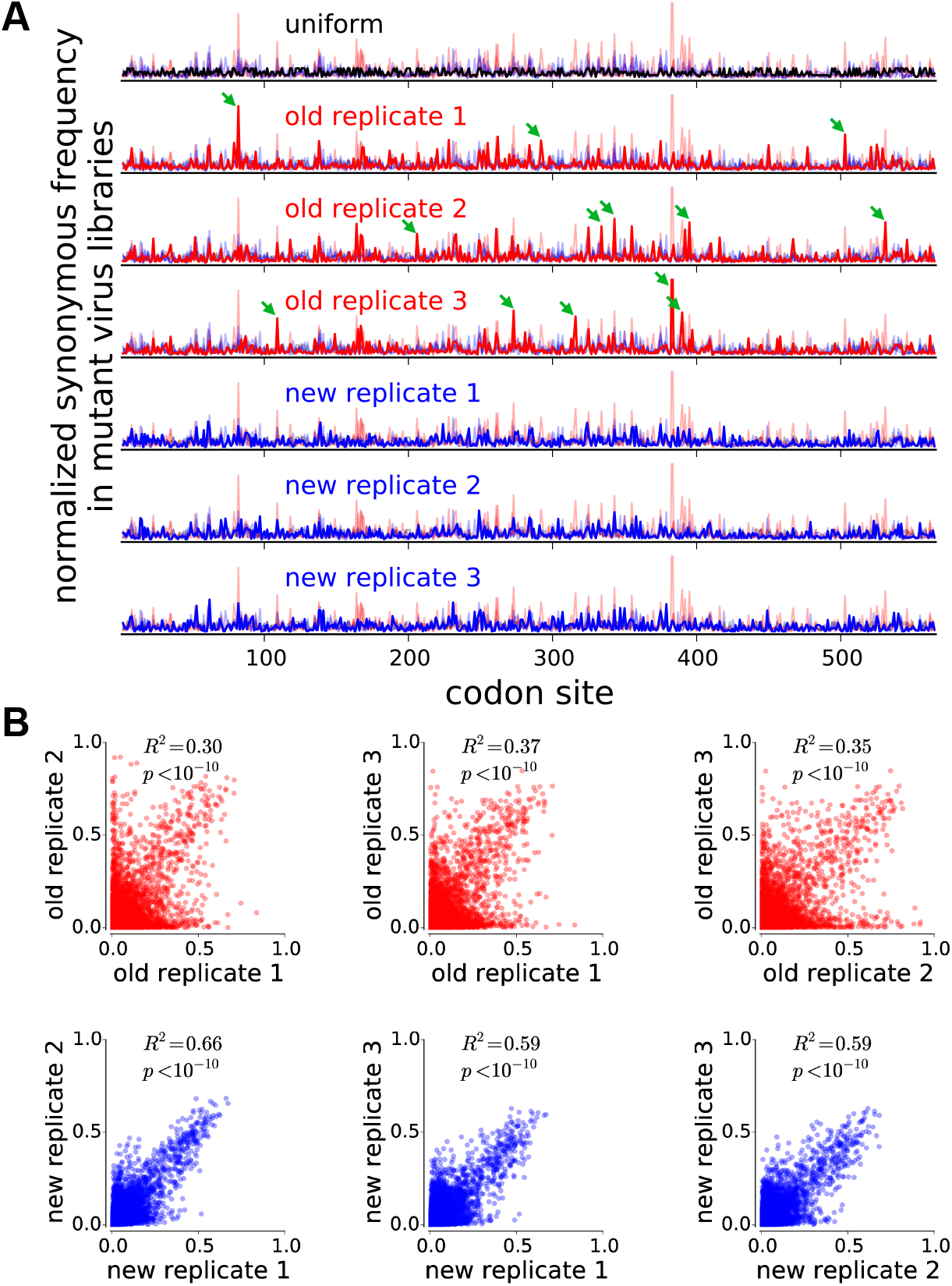
Our new experiments have increased reproducibility due to reduced bottlenecking during the generation of the mutant virus libraries. **(A)** Each row shows the synonymous mutation frequency for every site normalized to the total synonymous frequency for that sample. If synonymous mutations are sampled uniformly, the data should resemble the black line in the top row (the line is not completely straight because different codons have different numbers of synonymous variants). The next six rows show the synonymous mutation frequencies for each replicate of the old and new experiments, with that replicate shown with a thick line and the other replicates with thin lines. The old experiments have more bottlenecking as manifested by peaks at synonymous mutations present in viral variants that were stochastically enriched in that replicate (examples marked by green arrows). The differences between replicates are *not* due to differences in synonymous mutation frequencies in the plasmid libraries used to generate the viruses (Figure 2 - figure supplement 1). **(B)** The mutational effects measured in the new experiments are much more reproducible across replicates. Each plot shows the squared Pearson correlation coefficient for all site-specific amino-acid preferences measured in a pair of independent experimental replicates.

We next evaluated the reproducibility of our measurements of the effects of each amino-acid mutation. We estimated the effect of each mutation from its change in frequency in the mutant viruses relative to the original plasmid libraries, correcting for the site-specific error rates determined by sequencing unmutated virus and plasmid, and performing the analyses using the algorithms described in Bloom (2015) and implemented in the software at http://jbloomlab.github.io/dms_tools/. The results are quantified in terms of the *preference* of each site for each amino-acid; the set of all 20 preferences at a site can be thought of as representing the expected post-selection frequency of each amino acid at that site if all amino acids are initially present at equal frequencies.

Figure 2B shows the correlation between the amino-acid preferences from each experimental replicate. The replicate-to-replicate reproducibility is dramatically improved in our new experiments relative to those in Thyagarajan and Bloom (2014), with the average Pearson’s *R*^2^ increasing from 0.34 to 0.61. The new experiments are also largely free of the most problematic type of noise that plagued the original data, where an amino acid at a site is deemed highly preferred in one replicate but disfavored in another. Overall, these results demonstrate that our new strategies enable more reproducible measurement of the effects of all mutations to HA.

### The measurements better reflect the constraints on HA evolution in nature

We next tested whether our new measurements better describe the evolution of HA in nature. The accuracy with which experimental measurements of site-specific amino-acid preferences reflect the constraints shaping a protein’s evolution in nature can be quantified by comparing the phylogenetic fit of experimentally informed substitution models (Bloom, 2014a). We assembled a set of human and swine influenza HA sequences and fit substitution models using phydms (http://jbloomlab.github.io/phydms/; Bloom, 2016), which in turn uses `Bio++` (Gue´guen et al., 2013) for the likelihood calculations.

A substitution model informed by our new measurements described the natural evolution of HA better than a model informed by the older measurements from Thyagarajan and Bloom (2014), and vastly better than conventional non-site-specific substitution models (Table 1). Averaging the measurements from both studies improved phylogenetic fit even further, a finding consistent with previous work reporting that combining data from multiple deep mutational scanning studies of the same protein tends to improve substitution models (Doud et al., 2015).

**Table 1.** The site-specific amino-acid preferences measured in the new experiments offer an improved description of HA evolution in nature. Aikake information criterion (AIC) (Posada and Buckley, 2004) was used compare the maximum likelihood phylogenetic fit of several models to an alignment of seasonal human H1N1 and classical swine H1N1 HAs. The experimentally informed substitution models are of the form described in Bloom (2016) with the data from the average of all three replicates of the new or old experiments, or the average of the two. These models are compared to the variants of the substitution model of Goldman and Yang (1994) denoted as M0 and M8 in Yang et al. (2000) with the equilibrium codon frequencies estimated empirically using the F3X4 method. The best model is the one that combines all experimental data, but a model informed by the new experiments alone is better than one informed by the old experiments alone. To confirm that the experimentally informed models are superior because they are site specific, we fit a control model in which the experimental data is averaged across sites. The tree topology was fixed to that inferred by maximum likelihood using the M0 version of the Goldman-Yang model. The free parameters for each model were then optimized along with the branch lengths; optimized parameters are in the last column. The site-specific amino-acid preferences for the best model (new data + old data) are shown in Table 1 – figure supplement 1 and Table 1 – figure supplement 2; inferred differential selection between nature and the experiments is shown in Table 1 – figure supplement 3.

| model | $\Delta\text{AIC}$ | log likelihood | parameters (optimized + empirical):<br>optimized values |
| --- | --- | --- | --- |
| new data + old data | 0.0 | -14933.5 | 6 (6 + 0): $\beta = 1.82$, $\omega = 0.51$, $\kappa = 4.95$, $\phi_A = 0.40$, $\phi_C = 0.18$, $\phi_G = 0.20$ |
| new data | 197.6 | -15032.3 | 6 (6 + 0): $\beta = 1.80$, $\omega = 0.46$, $\kappa = 5.06$, $\phi_A = 0.40$, $\phi_C = 0.18$, $\phi_G = 0.20$ |
| old data | 341.2 | -15104.1 | 6 (6 + 0): $\beta = 1.40$, $\omega = 0.46$, $\kappa = 4.90$, $\phi_A = 0.39$, $\phi_C = 0.18$, $\phi_G = 0.20$ |
| Goldman-Yang M8 | 2156.8 | -16003.9 | 14 (5 + 9): $p_{\omega>1} = 0.01$, $\omega_{>1} = 1.91$, $p_\beta = 0.02$, $q_\beta = 0.76$, $\kappa = 4.94$ |
| new data + old data, averaged across sites | 2971.6 | -16419.3 | 6 (6 + 0): $\beta = 0.50$, $\omega = 0.20$, $\kappa = 5.38$, $\phi_A = 0.38$, $\phi_C = 0.18$, $\phi_G = 0.21$ |
| Goldman-Yang M0 | 2980.8 | -16418.9 | 11 (2 + 9): $\omega = 0.19$, $\kappa = 4.88$ |

The phylogenetic model fitting optimizes a parameter that accounts for differences in the stringency of selection between the experiments and natural evolution (Bloom, 2014b); a stringency parameter > 1 indicates that natural selection prefers the same amino acids as the experimental selections but with greater strength. The best model in Table 1 has a stringency parameter of 1.8. The site-specific amino-acid preferences for this model scaled by this stringency parameter are displayed in Table 1 – figure supplement 1 and Table 1 – figure supplement 2; text files with unscaled and scaled numerical values are in Supplementary file 1 and Supplementary file 2.

### A handful of sites are under very different selection in our experiments than in nature

We next asked whether there are sites in HA that evolve in nature in a way that is highly discordant with our experimental measurements. To do this, we used the approach in Bloom (2016) to identify selection in nature for amino acids that differ from the ones preferred in the deep mutational scanning. At most sites, the magnitude of differential selection is small (Table 1 – figure supplement 3), indicating that the experimentally measured preferences mostly parallel constraints on natural evolution. Sites that are under strong differential selection usually show conservative changes; for example, site 78 prefers isoleucine in nature but leucine in our deep mutational scanning.

One of the most striking exceptions to this general concordance between natural selection and our experiments can be given a clear explanation. At site 342 (328 in H3 numbering), the experimentally measured preference for tyrosine is at odds with nature’s strong preference for serine (Table 1 – figure supplement 3). The lab-adapted A/WSN/1933 strain used in our experiments differs from naturally occurring influenza in that it uses plasmin to cleave and activate HA (Lazarowitz et al., 1973; Goto and Kawaoka, 1998). Plasmin cleavage is enhanced by tyrosine at site 342 (Sun et al., 2010), so it is unsurprising that our experiments detected a preference at this site unique to the influenza strain we used. This example illustrates how the occasional deviations from the general concordance between deep mutational scanning experiments and natural selection can point to interesting biological mechanisms.

### Antigenic sites in HA’s globular head are highly tolerant of mutations, but stalk epitopes targeted by broadly neutralizing antibodies are not

We computed the inherent mutational tolerance of each site using the stringency-scaled amino-acid preferences from the combined datasets (Figure 3A). The mutational tolerance is mapped onto the structure of HA in Figure 3B.

**Figure 3.**
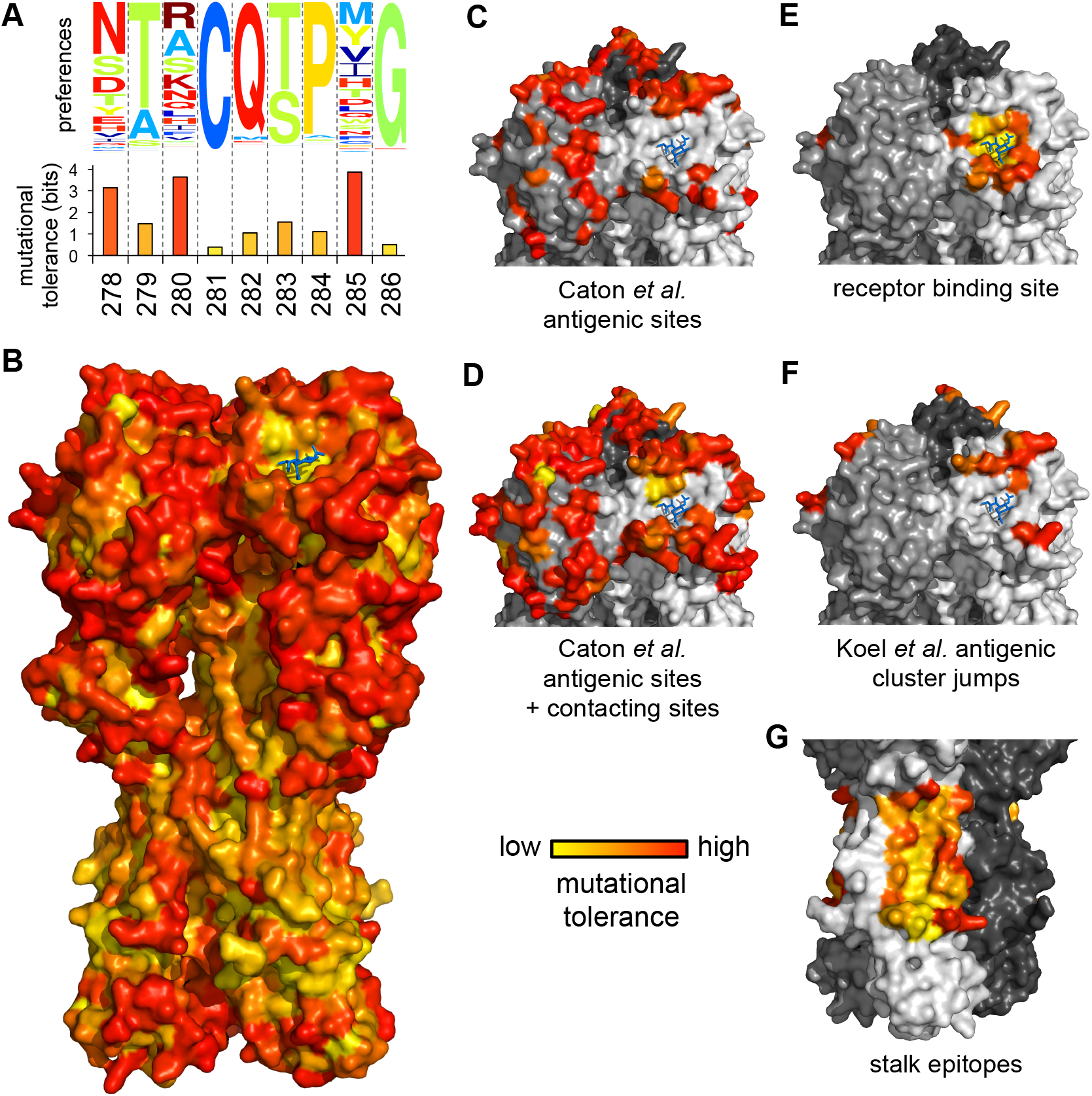
Antigenic sites in HA’s globular head have a high inherent tolerance for mutations, but HA’s stalk is relatively intolerant of mutations. **(A)** Mutational tolerance is calculated as the Shannon entropy of a site’s amino-acid preferences. **(B)** Mutational tolerance mapped onto the HA trimer (yellow indicates low tolerance, red indicates high tolerance, blue sticks show the sialic-acid receptor). **(C), (D)** The antigenic sites defined by Caton et al. (1982) have high mutational tolerance, as do the residues contacting these sites. **(E)** Conserved receptor-binding residues have low mutational tolerance. **(F)** Sites that contribute to antigenic duster jumps (Koel et al., 2013). **(G)** Sites in the footprints of four broadly neutralizing antibodies have low mutational tolerance. Shown are footprints of F10, CR6261, FI6v3, and CR9114 (Sui et al., 2009; Ekiert et al, 2009; Corti et al, 2011; Dreyfus et al, 2012). For panels B-H, tolerance is mapped onto PDB structure 1RVX (Gamblin et al., 2004). For panels C-G, each monomer is shown in a different shade of gray. Figure 3 - figure supplement 1 reports statistical analyses of whether subsets of sites have higher or lower tolerance than expected given their solvent accessibility. Figure 3 - figure supplement 2 shows the tolerance of the different domains of HA.

The H1 HA antigenic sites defined by Caton et al. (1982) are significantly more mutationally tolerant than the average site (Figure 3C), even after accounting for relative solvent accessibility (Figure 3 – figure supplement 1A). This high mutational tolerance extends to other solvent-exposed residues in contact with the antigenic sites (Figure 3D, Figure 3 – figure supplement 1B), indicating that the HA molecular surfaces commonly targeted by antibodies have a high inherent capacity for evolutionary change. This high mutational tolerance does not extend to the receptor-binding pocket (Figure 3E, Figure 3 – figure supplement 1C,D), but may be a feature of the sites that make the greatest contributions to the punctuated antigenic evolution of H3N2 and seasonal H1N1 HA (Koel et al., 2013) (Figure 3F), albeit not at a level that is statistically significant after correcting for solvent accessibility (Figure 3 – figure supplement 1E). These results support the findings of Thyagarajan and Bloom (2014) that the sites in HA that are the immunodominant targets of antibodies have a high inherent capacity to tolerate point mutations.

Perhaps in part because of the high mutational tolerance of the antigenic sites in its globular head, HA is adept at escaping antibody-mediated immunity (Smith et al., 2004; Bedford et al., 2014). New vaccines are being developed that aim to elicit immunity against other portions of HA (Krammer and Palese, 2013), most commonly regions in the stalk that are relatively conserved among naturally occurring strains. An important question is whether these stalk regions are conserved because they are inherently intolerant of point mutations, or simply because they are not currently under immune pressure. To answer this question, we examined the inherent mutational tolerance of the largely overlapping epitopes of four broadly neutralizing anti-stalk antibodies: F10 (Sui et al., 2009), CR6261 (Ekiert et al., 2009), FI6v3 (Corti et al., 2011), and CR9114 (Dreyfus et al., 2012). Visual inspection of Figure 3G shows that these stalk epitopes have a low mutational tolerance, a result that is confirmed by statistical analysis (Figure 3 – figure supplement 1F). Therefore, the epitopes that next-generation vaccines aim to target indeed have a reduced capacity for immune escape by point mutations.

We wondered if some of HA’s variation in mutational tolerance is explained by differences in the three ancient domains that compose the protein. HA is the product of a series of ancient insertions that merged a fusion domain, a receptor-binding domain, and a vestigial esterase domain (Rosenthal et al., 1998). We compared the inherent mutational tolerance of these three domains, again correcting for solvent accessibility. We found that sites in the receptor-binding domain have a significantly higher mutational tolerance than all sites in the protein, whereas sites in the fusion domain have a significantly lower mutational tolerance (Figure 3 – figure supplement 2). Therefore, HA’s antigenic evolvability is not just a consequence of the immunodominant antigenic sites themselves having high mutational tolerance, but also because these sites are found within a protein domain that is intrinsically more mutable than the rest of HA.

## Discussion

We have described new techniques that greatly improve the reproducibility of deep mutational scanning of influenza. The largest improvement appears to result from using a helper virus to generate virus mutant libraries without the bottlenecks that plague the creation of viruses purely from plasmids. We have used these techniques to more accurately measure the effects of all amino-acid mutations to HA. These measurements confirm our previous findings that HA’s propensity for immune escape is underpinned by the high inherent mutational tolerance of its antigenic sites. Importantly, our improved data also enable us to show that some regions of HA – including the stalk epitopes targeted by new broadly neutralizing antibodies – have a reduced capacity for evolutionary change. Overall, our work provides insight into how protein-intrinsic mutational tolerance shapes influenza evolution, and will provide a basis for using deep mutational scanning to improve quantitative models of viral evolution and understand virus-immune interactions.

## Materials and Methods

### Growth of HA-deficient helper virus in HA-expressing cells

MDCK-SIAT1-EF1a-WSN-HA cells were engineered to constitutively express the HA protein of A/WSN/1933 (H1N1) under control of the EF1a promoter by lentiviral transduction. HA surface expression was validated by flow cytometry (Figure 1 – figure supplement 1).

To generate HA-deficient helper virus, we seeded co-cultures of 293T cells (5 × 10^5^ cells per well) and MDCK-SIAT1-EF1a-WSN-HA cells (5×10^4^ cells cells per well) in 6-well dishes in D10 media (DMEM supplemented with 10% heat-inactivated FBS, 2 mM L-glutamine, 100 U of penicillin/ml, and 100 *µ*g of streptomycin/ml). After 24 hours, we transfected these co-cultures with bidirectional reverse-genetics plasmids for the seven non-HA segments of the A/WSN/1933 virus (pHW181-PB2, pHW182-PB1, pHW183-PA, pHW185-NP, pHW186-NA, pHW187-M, and pHW188-NS) (Hoffmann et al., 2000) plus a protein expression plasmid for WSN HA (pHAGE2-CMV-WSNHA, which importantly does *not* contain non-coding regions of the HA segment or a promoter for the transcription of negative-sense viral RNA). Transfection was performed with BioT transfection reagent (Bioland B01-02, Paramount, California) with each well receiving 250 ng of each plasmid. Twenty-two hours after transfection, we changed the media to WSN growth media (Opti-MEM supplemented with 0.5% heat-inactivated FBS, 0.3% BSA, 100 U of penicillin/ml, 100 *µ*g of streptomycin/ml, and 100 *µ*g of calcium chloride/ml). At 96 hours post-transfection, we passaged 400 *µ*l of the transfection supernatant to 15-cm dishes containing 4 × 10^6^ MDCK-SIAT1 cells (as a negative control) or MDCK-SIAT1-EF1a-WSN-HA cells in WSN growth media. HA-deficient helper virus could only be propagated in the HA-expressing cells as expected (Figure 1 – figure supplement 1) We collected the expanded helper virus from these cells after 68 hours, aliquoted, and froze aliquots at −80°C. We titered the helper virus in MDCK-SIAT1-EF1a-WSN-HA cells by TCID_50_. We obtained titers between 10^3^ and 10^4^ TCID_50_ per *µ*l when titering in MDCK-SIAT1-EF1a-WSN-HA cells, and no cytopathic effect except with extremely concentrated helper virus in MDCK-SIAT1 cells (Figure 1 – figure supplement 1).

### HA plasmid mutant libraries

Codon mutagenesis was performed as described in Thyagarajan and Bloom (2014) except that we performed one overall round of the PCR mutagenesis to yield a lower mutation rate (Figure 1 – figure supplement 2). Ligation and eletroporation were also performed as in Thyagarajan and Bloom (2014), except that we cloned the inserts into both pHW2000 (Hoffmann et al., 2000) and pHH21 (Neumann et al., 1999) plasmid backbones. All steps were performed in triplicate. For each replicate, we pooled over 3 million transformants, cultured in LB for 3 hours in shaking flasks at 37^o^C, and maxi-prepped plasmid libraries.

### Generation of mutant HA virus libraries from mutant plasmids and helper viruses

To generate mutant virus libraries, we transfected 293T cells with a DNA mixture containing one of the three pHH21-MutantHA libraries (or the wild-type pHH21-WSN-HA control) and protein expression plasmids for the four proteins that compose the ribonucleoprotein complex, using plasmids HDM-Nan95-PA, HDM-Nan95-PB1, HDM-Nan95-PB2, and HDM-Aichi68-NP (Gong et al., 2013). Specifically, we plated 293T cells in D10 at a density of 8 × 10^5^ per well in 6-well plates, changed the media to fresh D10 after 16 hours, and then four hours later transfected cells with 500 ng of the HA reverse-genetics plasmid plus 375 ng of each of the PA, PB1, PB2, and NP plasmids using BioT. Twenty-four hours after transfection, we infected the cells with HA-deficient helper virus by making an inoculum of 1.3×10^3^ TCID_50_ per *µ*l in WSN growth media, aspirating the D10 media from the cells, and adding 2 ml of inoculum to each well. After 3 hours, we removed the inoculum by aspiration and added 2 ml of WSN growth media supplemented with 5% D10. Twenty-four hours after helper virus infection, we collected the supernatants for each replicate, stored aliquots at −80°C, and titered in MDCK-SIAT1 cells. Of note, we found that helper viruses that had been passaged more than once in MDCK-SIAT1-EF1a-WSN-HA cells tended to become less effective at rescuing fully replication competent viruses following infection of transfected cells, so we exclusively used single-passage helper virus in these experiments.

We passaged these transfection supernatants to create a genotype-phenotype link and impose functional selection on HA. We passaged over 9 × 10^5^ TCID_50_ at an MOI of 0.0075 TCID_50_ per cell. Specifically, for each library, we plated ten 15-cm dishes with 6 × 10^6^ MDCK-SIAT1 cells per dish and allowed cells to grow for 20 hours, at which point they had reached a density ∼ 1.25 × 10^7^ cells per dish. We then replaced the media in each dish with 25 ml of WSN growth media in each dish containing 3.7 TCID_50_ of virus per *µ*l. We allowed virus replication to proceed for 40 hours before collecting virus from the supernatant for sequencing.

### Barcoded subamplicon sequencing

For each of the three replicate HA virus libraries and the wild-type HA virus, we extracted viral RNA by ultracentrifuging 24 ml of supernatant at 22,000 rpm in a Beckman Coulter SW28 rotor. RNA was extracted using the Qiagen RNeasy kit by resuspending the viral pellet in 400 *µ*l buffer RLT freshly supplemented with β-mercaptoethanol, pipetting 30 times, transferring to an RNase-free microcentrifugefuge tube, adding 600 *µ*l freshly-made 70% ethanol, and continuing with the manufacturer’s recommended protocol, eluting the final RNA product in 40 *µ*l of RNase-free water. HA was then reverse transcribed using AccuScript Reverse Transcriptase (Agilent 200820) with the primers WSNHA-For (5’-AGCAAAAGCAGGGGAAAATAAAAACAAC-3’) and WSNHA-Rev (5’-AGTAGAAACAAGGGTGTTTTTCCTTATATTTCTG-3’).

We generated PCR amplicons of HA for each of the eight samples (three replicate plasmid DNA libraries, three corresponding virus libraries, one wild-type plasmid DNA, and one wild-type virus) using KOD Hot Start Master Mix (71842, EMD Millipore) with the PCR reaction mixture and cycling conditions described in Bloom (2014a) and the primers WSNHA-For and WSNHA-Rev. The templates for these reactions were 2 *µ*l of cDNA (for the virus-derived samples) or 2 *µ*l of plasmid DNA at 10ng/*µ*l. To ensure that the number of molecules used as template did not bottleneck diversity, parallel PCR reactions were run with a standard curve of template molecules and all products were analyzed by band intensity after agarose gel electrophoresis; all samples used ≥ 10^6^ molecules as template for PCR. We purified these PCR amplicons using Agencourt AMPure XP beads (bead-to-sample ratio 0.9) (Beckman Coulter).

These PCR amplicons were quantified using Quant-iT PicoGreen dsDNA Assay Kit (Life Technologies) and used as the templates for the barcoded-subamplicon sequencing in Figure 1B. We performed the first round of PCR (“PCR 1”) in six parallel reactions (one for each of the six HA subamplicons) for each of the eight samples. Each reaction contained 12 *µ*l 2X KOD Hot Start Master Mix, 2 *µ*l forward primer diluted to 5*µ*M, 2 *µ*l reverse primer diluted to 5*µ*M, and 8 *µ*l purified amplicon diluted to 0.5ng/*µ*l (primer sequences for PCR 1 and PCR 2 are provided in Supplementary file 4). In addition to containing sequences targeting regions in HA, the forward and reverse primers for PCR 1 each contain an 8-base degenerate barcode and partial Illumina sequencing adaptors. To limit the generation of PCR artifacts, we performed only 9 cycles of PCR for PCR 1 using the following program: 1. 95^o^C for 2:00, 2. 95^o^C for 0:20, 3. 70^o^C for 0:01, 4. 54^o^C for 0:20, 5. 70^o^C for 0:20, 6. Go to 2 (8 times), 7. 95^o^C for 1:00, 8. 4^o^C hold. The denaturation step after cycling ensures that identical barcode pairs are not annealed at the end, so that most double-stranded molecules entering PCR 2 will contain two unique barcoded mutants. PCR 1 products were purified by Ampure XP (bead-to-sample ratio 1.0), quantified with Quant-iT PicoGreen, and diluted to 0.5 ng/*µ*l.

We then mixed all six subamplicons from each experimental sample at equal concentrations and diluted these subamplicon pools such that the number of template molecules used in PCR 2 was less than the anticipated sequencing depth to ensure multiple reads per barcode. Specifically, we reduced the total amount of DNA for each experimental sample used as template in PCR 2 to 9.24 × 10^-4^ ng, which corresponds to 1.54 × 10^-4^ ng of each of the six subamplicons, corresponding to approximately 3.5 × 10^5^ double-stranded DNA molecules (or 7 × 10^5^ uniquely-barcoded single-stranded variants) per subamplicon per sample.

We performed PCR 2 for each sample with the following reaction conditions: 20 ul 2X KOD Hot Start Master Mix, 4 *µ*l forward primer UniversalRnd2for diluted to 5 *µ*M, 4 *µ*l reverse primer indexXXRnd2rev diluted to 5 *µ*M (a different index for each experimental sample), 9.24 × 10^-4^ ng of the subamplicon pool of PCR 1 products described above, in a total volume of 40 *µ*l. We used the following thermal cycling program: 1. 95^o^C for 2:00, 2. 95^o^C for 0:20, 3. 70^o^C for 0:01, 4. 55^o^C for 0:20, 5. 70^o^C for 0:20, 6. Go to 2 (23 times), 7. 4^o^C hold. PCR 2 products were purified by Ampure XP (bead-to-sample ratio 1.0), quantified with Quant-iT PicoGreen, and equal amounts of each experimental sample were mixed and purified by agarose gel electrophoresis, excising the predominant DNA species at the expected size of approximately 470 bp. Sequencing was performed on one lane of a flow cell of an Illumina HiSeq 2500 using 2×250 bp paired-end reads in rapid-run mode.

### Inference of amino-acid preferences from sequencing data

We used dms tools (http://jbloomlab.github.io/dms_tools/), version 1.1.12, to align subamplicon reads to a reference HA sequence, group barcodes to build consensus sequences, quantify mutation counts at every site in the gene for each experimental sample, and infer site-specific amino-acid preferences based on mutation frequencies pre- and post-selection using the algorithm described in Bloom (2015). The code that performs these analyses is in Supplementary file 3.

### Phylogenetic modeling using amino-acid preferences

We sub-sampled human and swine H1 sequences (1 sequence per host per year) from the set of sequences from Thyagarajan and Bloom (2014), removed identical sequences, and built a sequence alignment. We then used phydms version 1.1.0 (http://jbloomlab.github.io/phydms/; Bloom, 2016), which in turn uses `Bio++` (Gue´guen et al., 2013) for the likelihood calculations, to compare experimentally informed codon substitution models and other non-site-specific substitution models. The code that performs these analyses is in Supplementary file 3.

### Statistical tests

Multiple linear regression of the continuous dependent variable of site entropy as a function of the continuous independent variable of relative solvent accessibility and a binary indicator of a site belonging to a specific classification (e.g. “antigenic sites”) was performed with the same classifications as described in Thyagarajan and Bloom (2014). Additional classifications were obtained from Koel et al. (2013) for sites responsible for antigenic cluster transitions in H3N2 and seasonal H1N1 (sites 158, 168, 169, 171, 172, 202, and 206 in sequential WSN H1 numbering starting with the initiating methionine), and the sites within antibody footprints of broadly-neutralizing antibodies F10, CR6261, FI6v3, and CR9114 (sites 25, 45, 46, 47, 48, 49, 305, 306, 307, 332, 361, 362, 363, 364, 379, 381, 382, 384, 385, 386, 388, 389, 391, 392, 395, 396, 399, and 400 in sequential WSN H1 numbering starting with the initiating methionine) (Sui et al., 2009; Ekiert et al., 2009; Corti et al., 2011; Dreyfus et al., 2012). Definition of the protein domains within HA were from Gamblin et al. (2004) (HA1 fusion domain: 18-72, 291-340; HA1 vestigial esterase domain: 73-125, 279-290; HA1 receptor binding domain: 126-278; HA2 fusion domain: 344-503; all sites in sequential H1 numbering starting with the initiating methionine). The code that performs these analyses is in Supplementary file 3.

### Availability of data and computer code

Sequencing data are available from the Sequence Read Archive under accession numbers SRR3113656 (mutant DNA library 1), SRR3113657 (mutant DNA library 2), SRR3113658 (mutant DNA library 3), SRR3113660 (mutant virus library 1), SRR3113661 (mutant virus library 2), SRR3113662 (mutant virus library 3), SRR3113655 (wild-type DNA control), and SRR3113659 (wild-type virus control). An iPython notebook (and a static HTML version of it) for all analyses is in Supplementary file 3. A Python script for visualizing mutational tolerance on the HA structure in PyMol is in Figure 3 – source code 1.

## Acknowledgments

We thank Bargavi Thyagarajan for performing the PCR mutagenesis of the HA gene. We thank Anice Lowen for discussions that helped inspire the idea of using a helper virus to generate the mutant virus libraries. This work was supported by the NIGMS of the NIH under grant R01 GM102198. M.B.D. was supported in part by a fellowship from the Seattle Chapter of the Achievement Rewards for College Scientists Foundation.

---

**Figure 1 – figure supplement 1:**
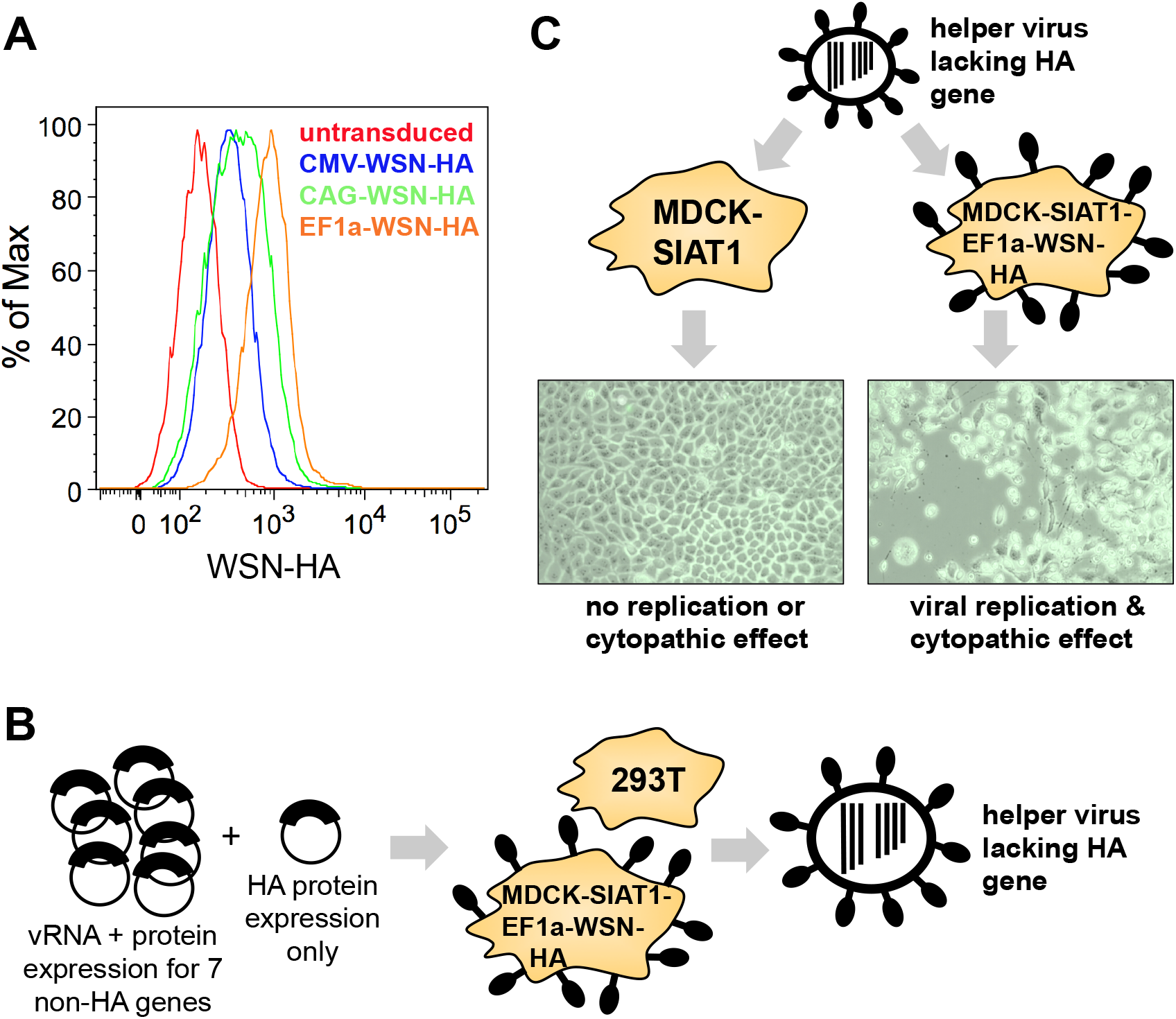
An HA-deficient helper virus can replicate in cells constitutively expressing HA protein. **(A)** We engineered MDCK-SIAT1 cells by lentiviral transduction to constitutively express the HA protein of the A/WSN/1933 strain under control of the EF1a promoter (MDCK-SIATl-EFla-WSN-HA cells), which provided higher expression levels than the CMV or CAG promoters. Transduced and untransduced cells were stained with a 1:100 dilution of mouse polyclonal anti-WSN serum, followed by a 1:100 dilution of APC-conjugated anti-mouse IgG for secondary staining. **(B)** We transfected a co-culture of these MDCK-SIAT1-EF1a-WSN-HA and 293T cells with bidirectional reverse-genetics plasmids (Hoffmann et al., 2000) for the seven non-HA segments of A/WSN/1933 plus a protein expression plasmid for HA. **(C)** The resulting transfection supernatant contained HA-deficient helper virus that could be propagated in MDCK-SIATl-EFla-WSN-HA cells but *not* in standard MDCK-SIAT1 cells. This virus typically reached titers of ~ 10^3^ TCID_50_ per *μ*l when titered on the MDCK-SIATl-EFla-WSN-HA cells. This titer was about 3-fold lower than that obtained if we included an HA GFP segment similar to that described in Marsh et al. (2007) that contains GFP flanked by the noncoding and 80 coding nucleotides. This difference in titer between an HA-deficient and HA-GFP virus is comparable to that reported by Marsh et al. (2007).

**Figure 1 – figure supplement 2:**
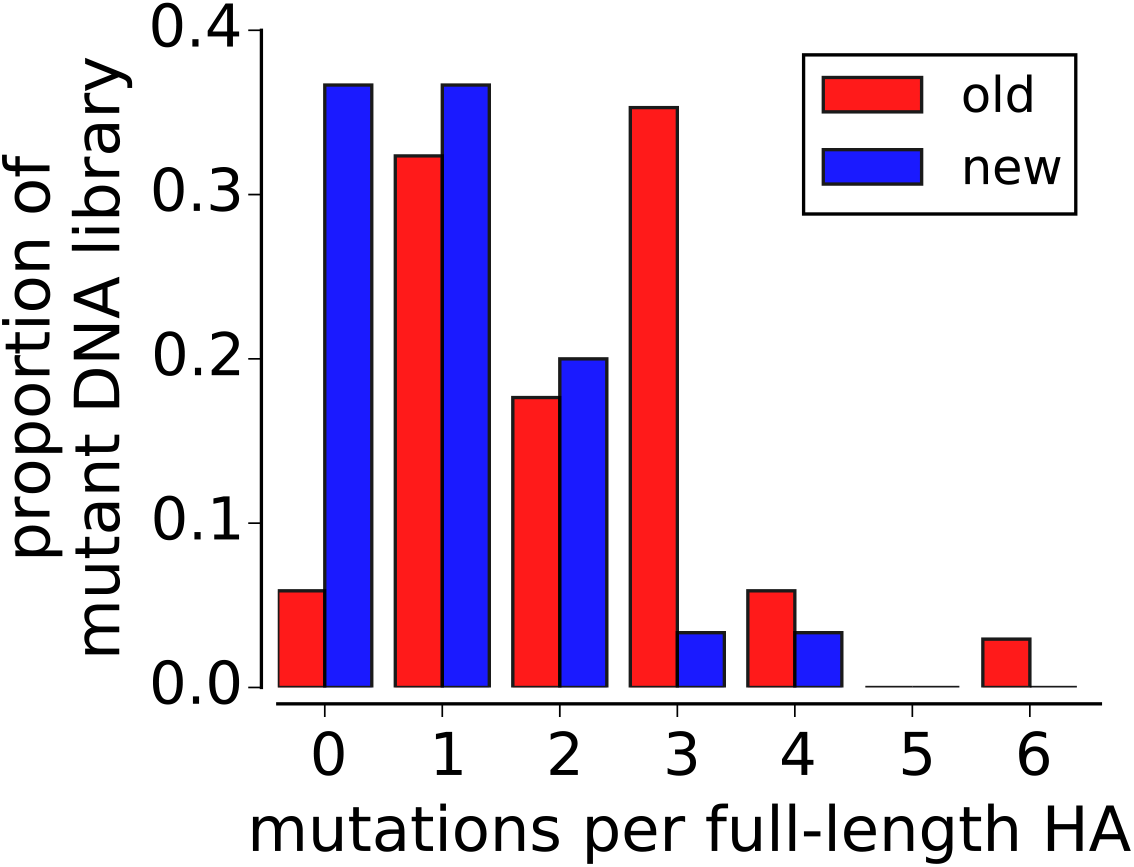
The mutant plasmid DNA library used in this study (“new”) has a lower mutation rate than the library used by Thyagarajan and Bloom (2014) (“old”). The old library was generated using two rounds of codon mutagenesis, leading to an average of two mutations per HA; the new library used only one round of mutagenesis, resulting in an average of one mutation per HA. At least 30 clones of each library were Sanger sequenced across the entire gene.

**Figure 1 – figure supplement 3:**
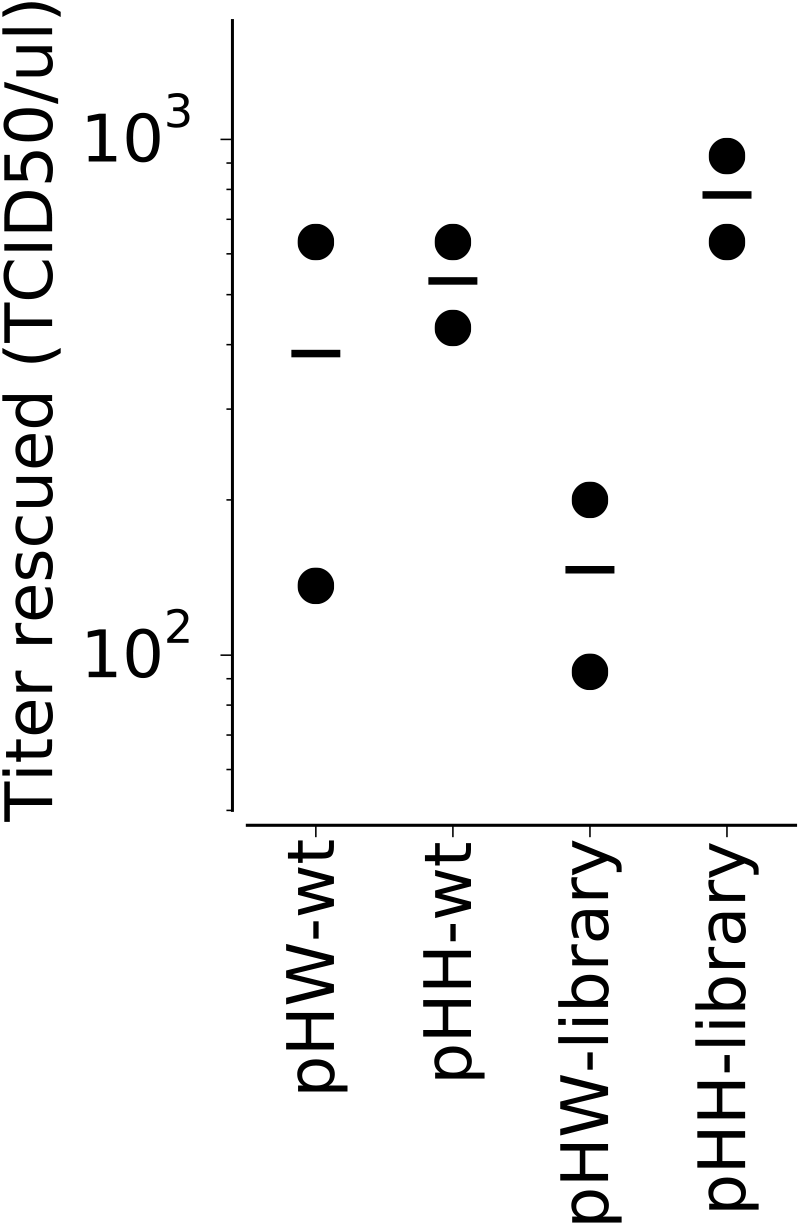
Mutant virus library generation is more efficient when HA is encoded on the pHH21 plasmid. The pHH21 plasmid contains a single RNA polymerase I promoter for transcription of negative sense HA vRNA (Neumann et al., 1999); this vRNA molecule must then associate with the viral polymerase complex for mRNA transcription and protein expression for proper virus generation. The pHW2000 plasmid is similar to pHH21, but also contains an RNA polymerase II promoter (Hoffmann et al., 2000) for the transcription of both negative sense HA vRNA and positive sense HA mRNA directly off the plasmid, so that HA protein expression is not limited by the transcription of mRNA by the viral polymerase. Cells were transfected with the indicated plasmid containing wild-type or mutant library HA along with protein expression plasmids for the viral polymerase-related proteins (PB2, PB1, PA, and NP). After 24 hours, cells were infected with helper virus, and 24 hours after infection, cell supernatants were titered in MDCK-SIAT1 cells to quantify the amount of virus generated. We hypothesize that the lower titer when using the pHW plasmid is due to expression of more HA mutants per cell, some of which might act as dominant negatives. Each virus generation was performed in duplicate, with a bar marking the mean of the two experiments.

**Figure 1 – figure supplement 4:**
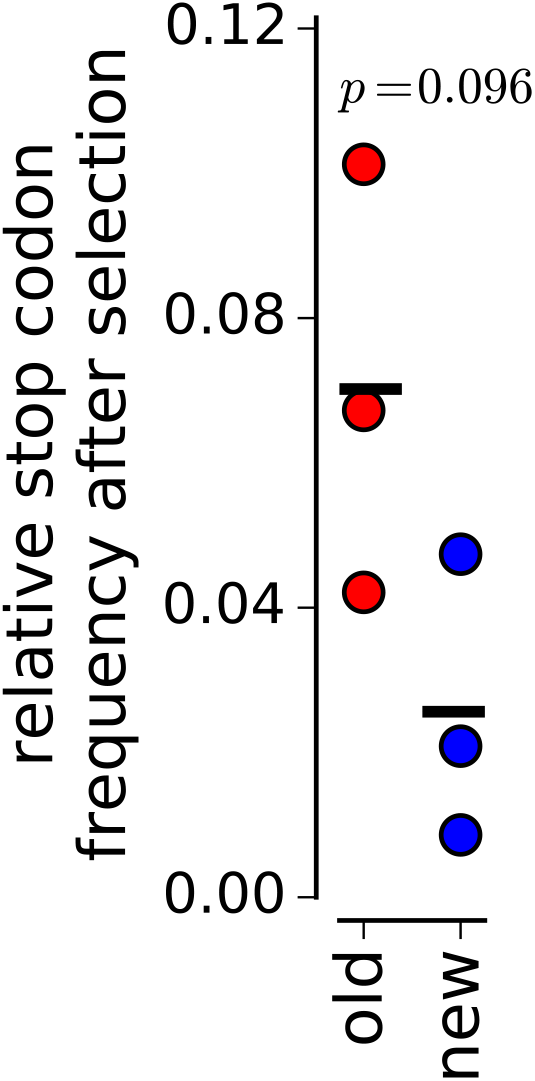
Purging of stop codons is more complete in our new experiments than in the previous one. Each point shows the relative fraction of stop codons remaining after selection for one of the three replicates. The stop codon frequencies in the wild-type plasmid and virus samples are subtracted from the mutant plasmid and mutant virus samples to correct for errors arising during sample preparation and sequencing. “Old” refers to libraries from Thyagarajan and Bloom (2014); “new” refers to libraries in the current study. We hypothesize that the stronger selection against stop codons in the new experiments is the result of better genotype-phenotype linkage imposed by the the lower MOI used for viral passage, which leads to more effective selection on each mutation. Bars show the means for each set of libraries; the p-value was calculated with a two-sided T-test.

**Figure 2 – figure supplement 1:**
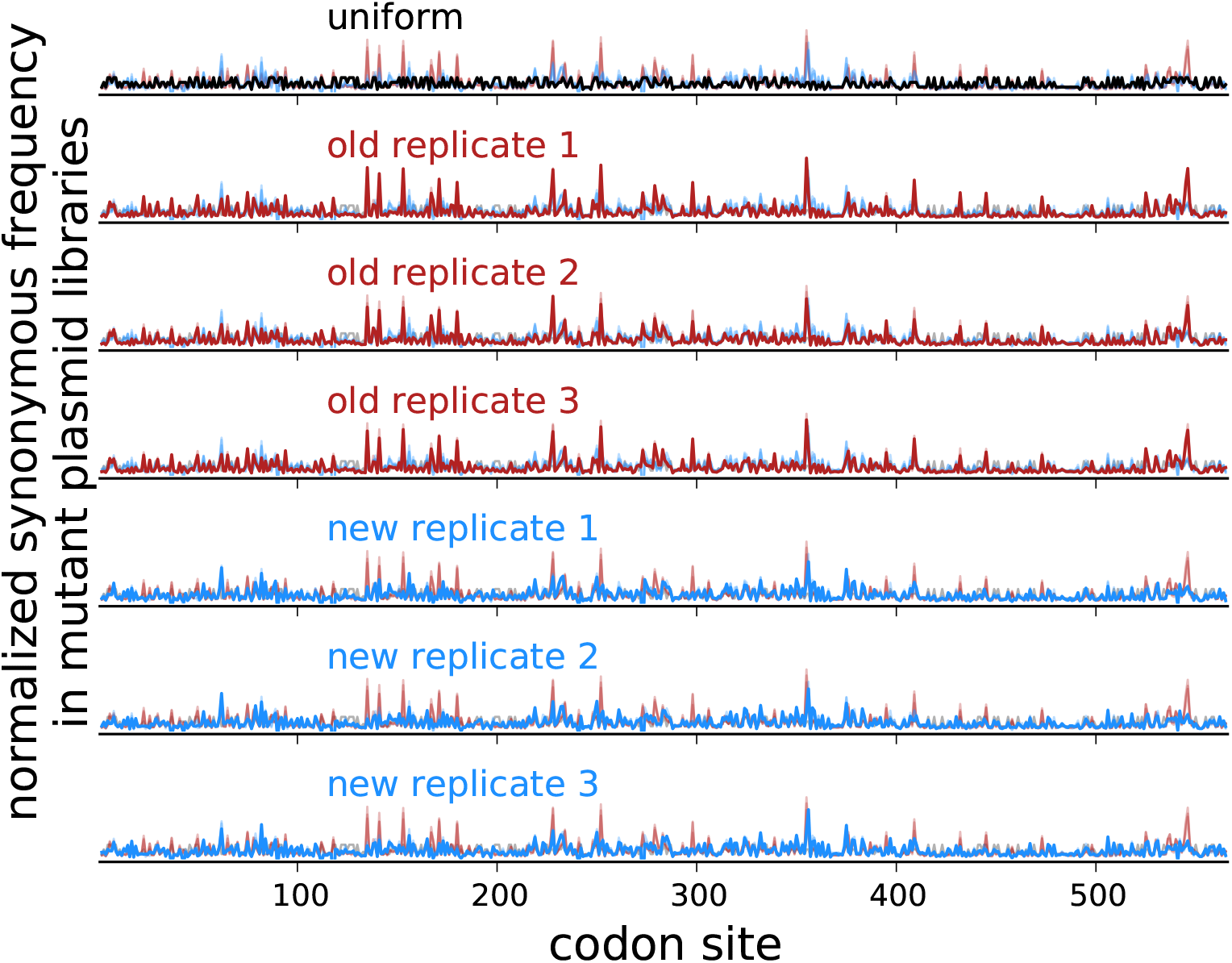
Synonymous frequency peaks observed in bottlenecked virus libraries are not due to the composition of plasmid mutant libraries. Shown for each replicate is the normalized synonymous mutation frequency for plasmid mutant libraries in the same form as Figure 2. The frequency of synonymous mutations in the plasmid mutant libraries is highly reproducible among replicates. Although there are some sites with peaked frequencies (likely due to PCR biases during mutagenesis), these sites are consistent across replicates and do not correspond to the peaks in the mutant virus libraries shown in Figure 2A.

**Figure 3 – figure supplement 1:**
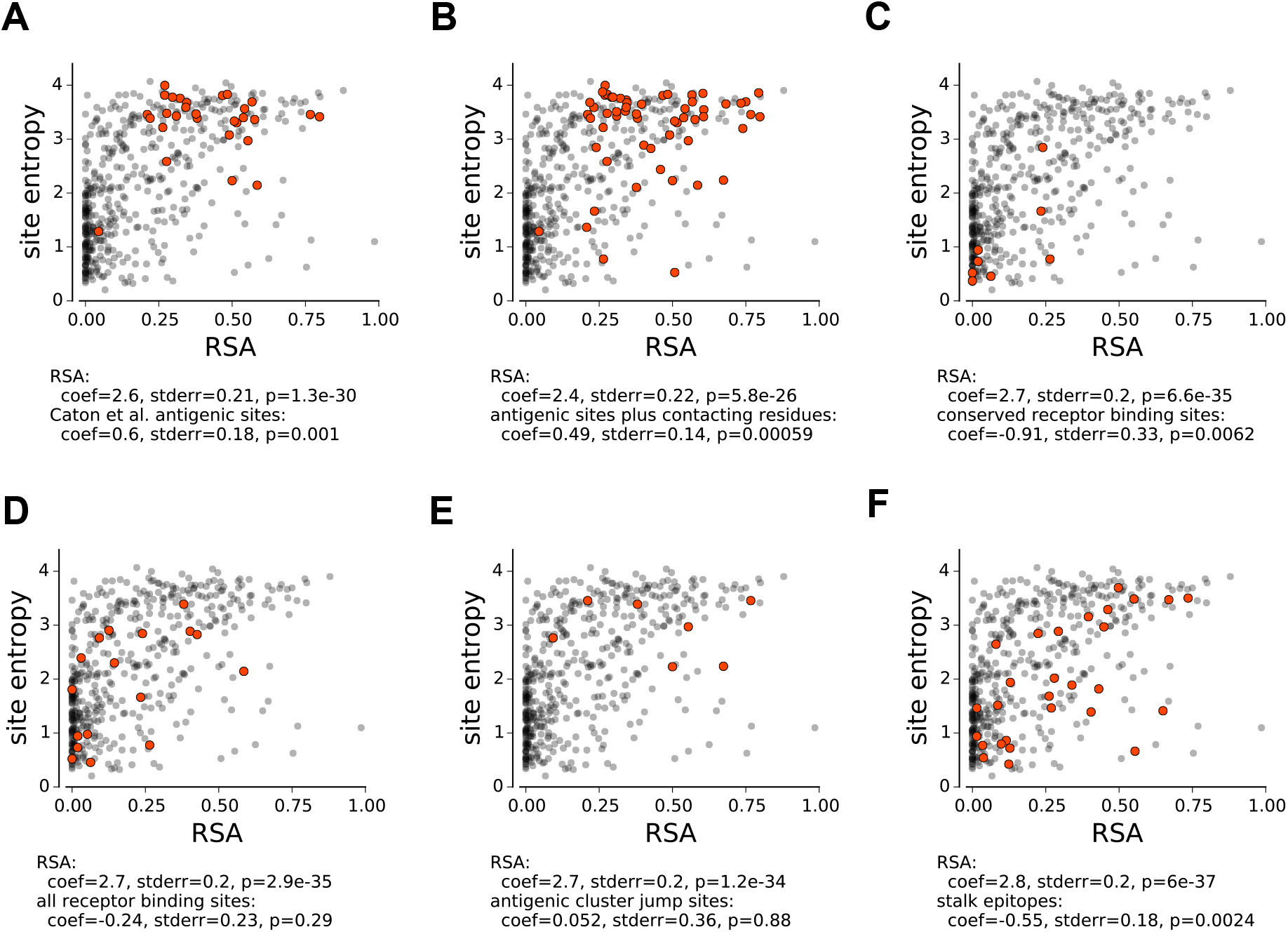
Statistical analyses of whether sets of sites have higher or lower mutational tolerance than expected given their solvent accessibility. For each panel, a group of sites is selected and the results are shown for multiple linear regression of site entropy as a function of relative solvent accessibility and whether or not the site belongs to that group. **(A)** Antigenic sites defined by Caton et al. (1982) and **(B)** these sites plus their contacts have significantly higher mutational tolerance than expected from their solvent accessibility. **(C)** Conserved receptor binding sites have significantly lower mutational tolerance. **(D)** All sites contacting receptor have typical mutational tolerance. **(E)** Antigenic cluster jump sites defined by Koel et al. (2013) have typical mutational tolerance. **(F)** Sites in the overlapping footprints of broadly neutralizing antibodies F10, CR6261, FI6v3, and CR9114 have significantly lower mutational tolerance.

**Figure 3 – figure supplement 2:**
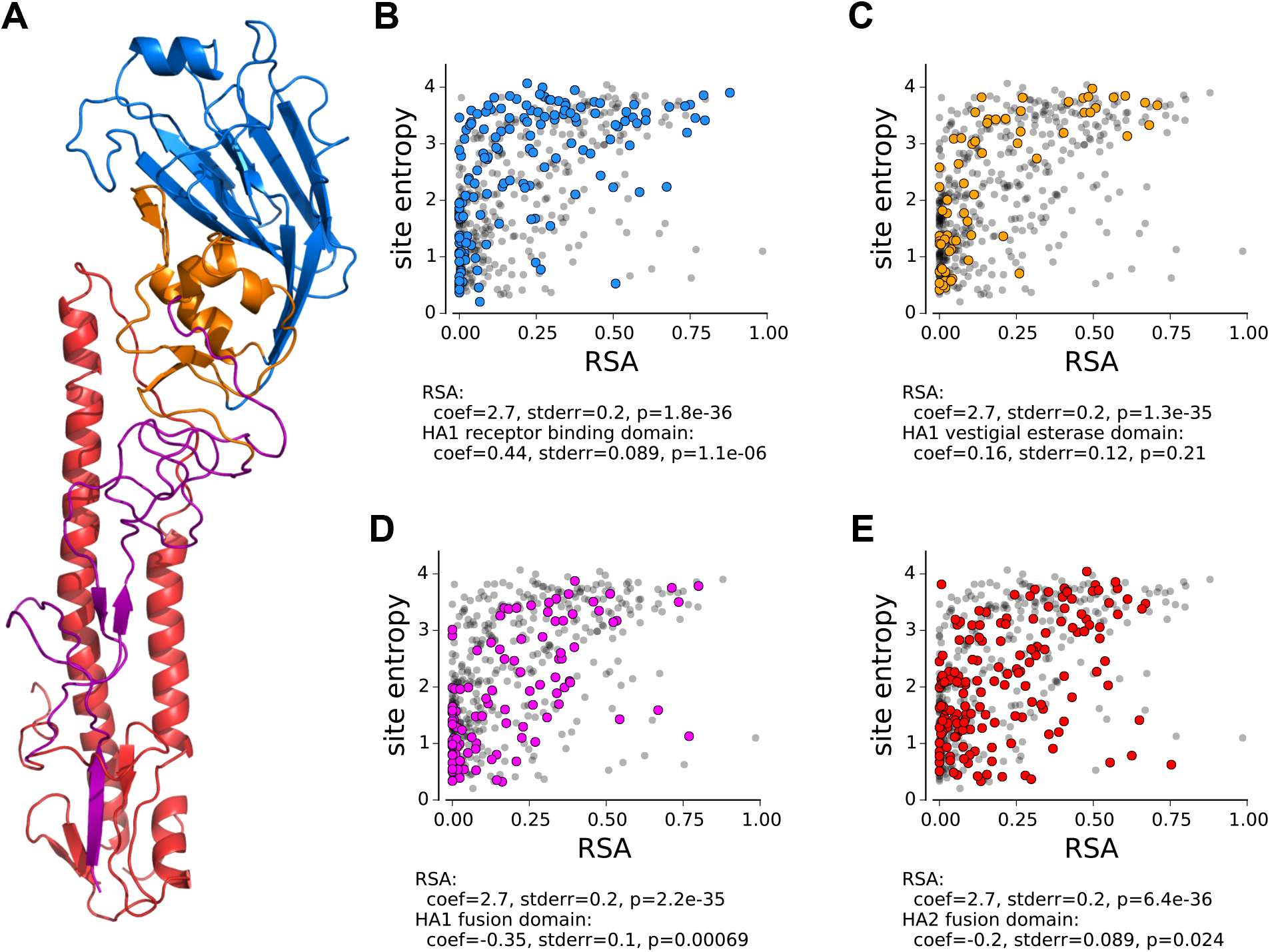
The mutational tolerance of HA’s domains. **(A)** The domain architecture of HA. The receptor-binding domain is blue, the vestigial esterase domain is orange, and the fusion subdomains of HA1 and HA2 are purple and red, respectively. **(B)** The mutational tolerance of the receptor-binding domain is significantly higher than the rest of the protein. **(C)** The mutational tolerance of the vestigial esterase domain is not significantly different than the rest of the protein. **(D), (E)** The mutational tolerance of the fusion subdomains of HA1 and HA2 is significantly lower than the rest of the protein. Significance is assessed using multiple linear regression, correcting for solvent accessibility as in Figure 3 – figure supplement 1.

**Figure 3 – source code 1**: PyMol script for visualization of mutational tolerance on the HA crystal structure.

**Table 1 – figure supplement 1:**
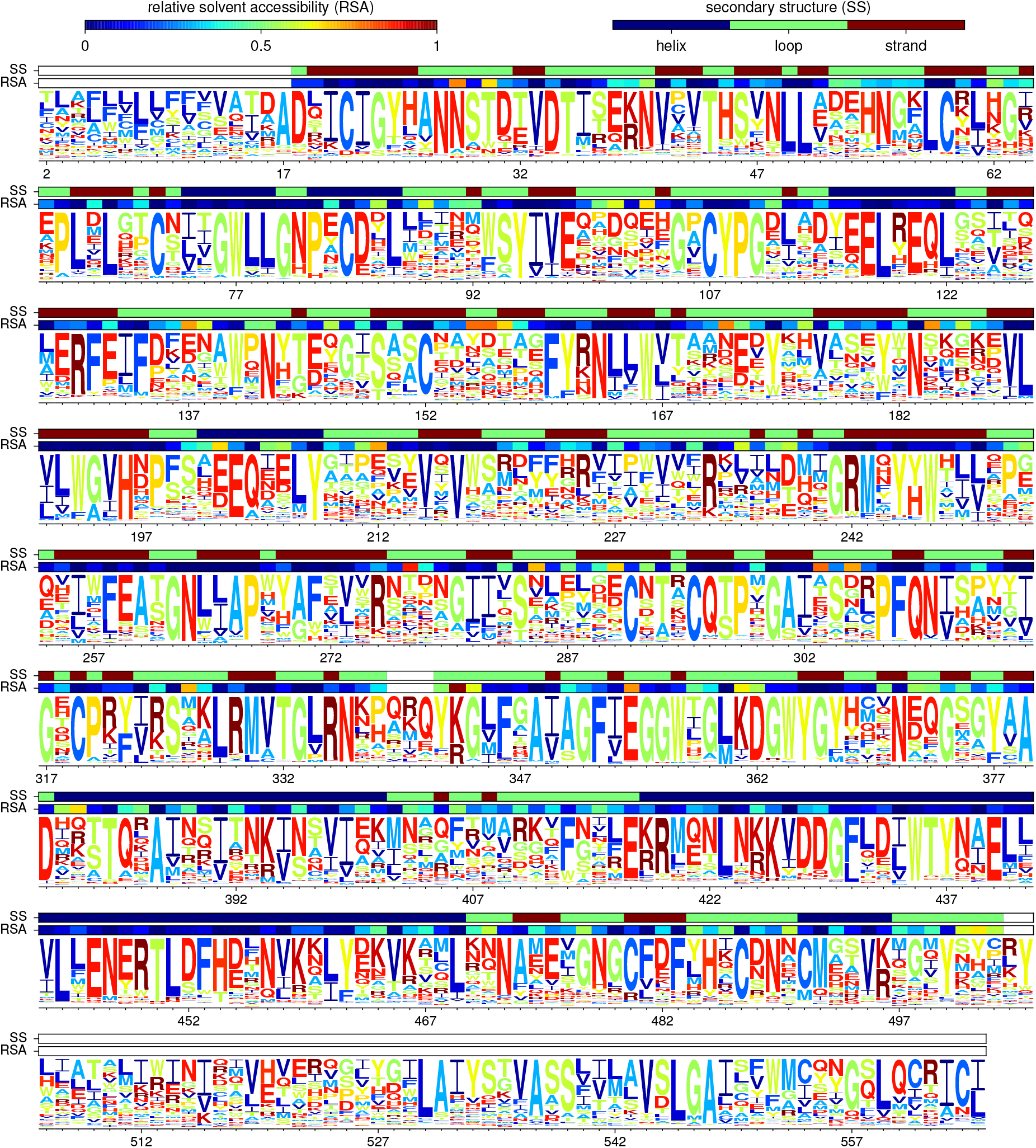
The site-specific amino-acid for the combined new data + old data scaled by the stringency parameter inferred in Table 1. Residues are numbered according to sequential numbering of the A/WSN/1933 HA beginning with 1 at the N-terminal methionine. This first residue is not shown since it was not mutagenized in the deep mutational scanning.

**Table 1 – figure supplement 2:**
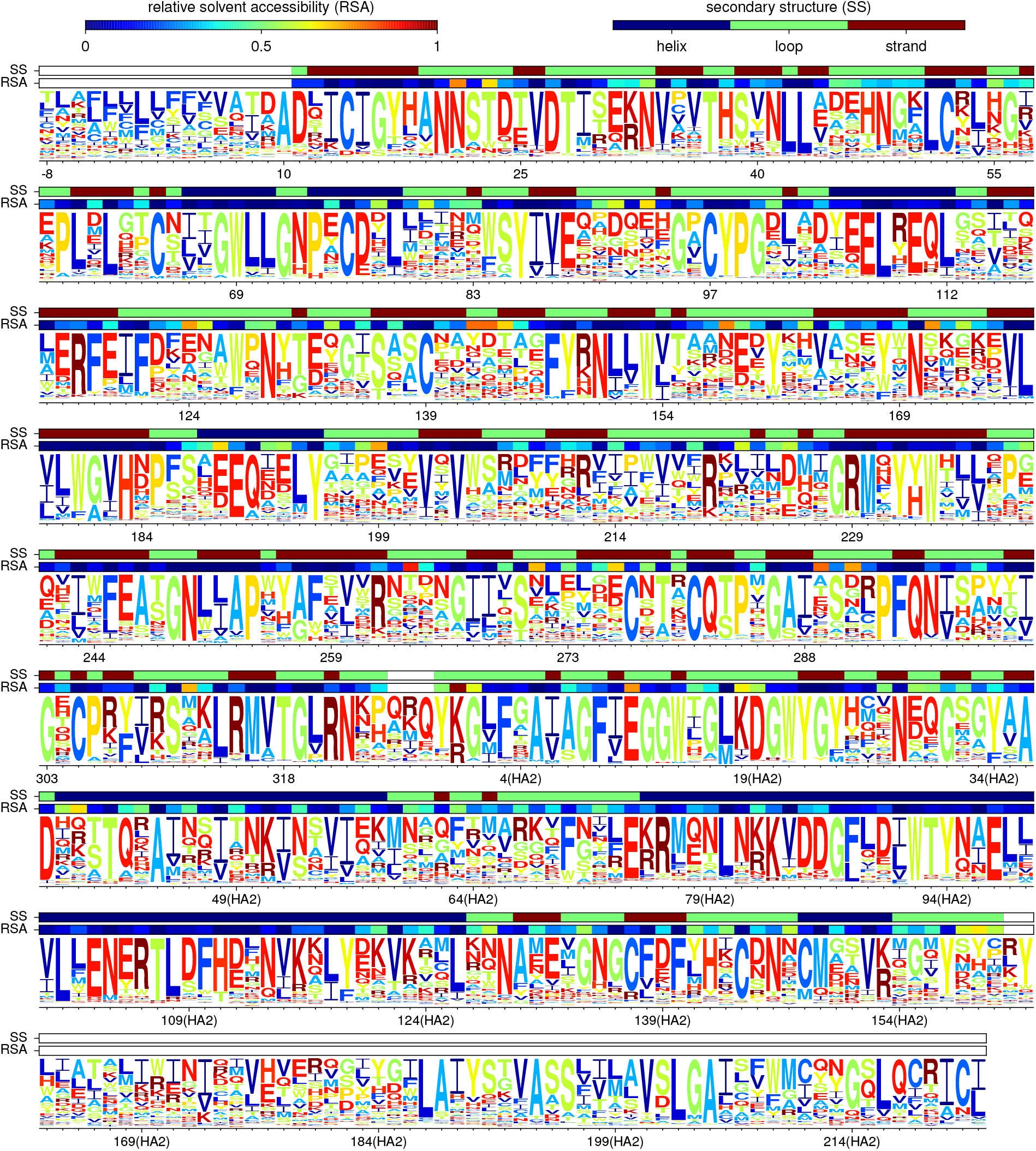
This figure is like Table 1 - figure supplement 1 except that the residues are numbered using the H3 numbering scheme.

**Table 1 – figure supplement 3:**
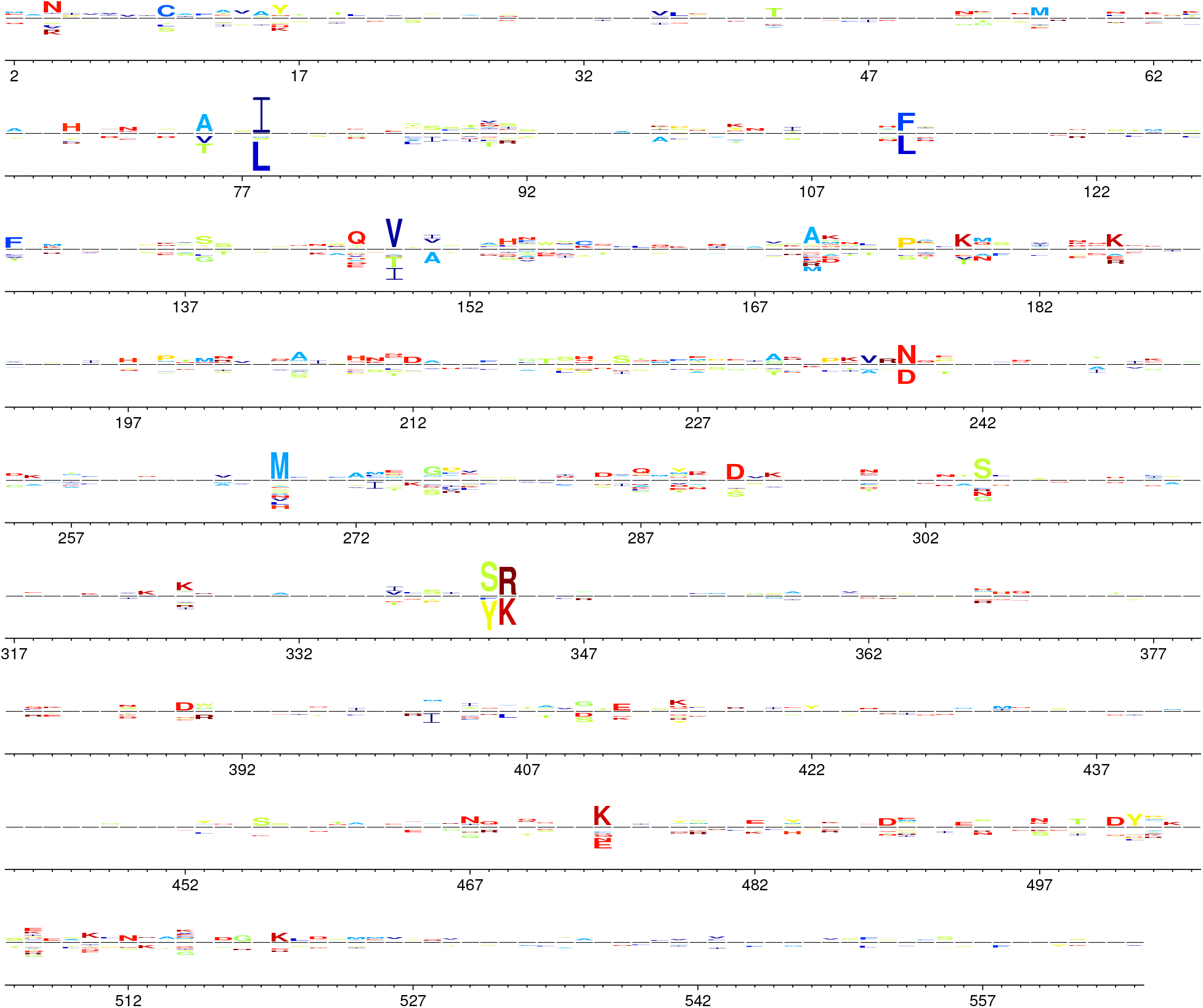
Differential selection between the deep mutational scanning and natural evolution for the alignment of human seasonal H1N1 and classical swine H1N1 HAs. At each site, the height of a letter above or below the center line indicates that differential selection for or against that amino acid. The differential selection is inferred using the approach described in Bloom (2016). At most sites, the differential selection is very small, showing that the experimental measurements are mostly concordant with natural selection on HA. The residues are numbered according to sequential numbering of the A/WSN/1933 HA beginning with 1 at the N-terminal methionine.

**Supplementary file 1**: Text file with the overall merged site-specific amino-acid preferences (average of the new and old data).

**Supplementary file 2**: Text file with the overall merged site-specific amino-acid preferences (average of the new and old data) scaled by the stringency parameter from Table 1.

**Supplementary file 3**: A ZIP file containing the data and code for all data analysis and figure generation. The analysis is performed by the iPython notebook in this file.

**Supplementary file 4**: Text file with primer sequences for the barcoded subamplicon sequencing.

